# Diminishing Arousal Shifts the Balance of Cortical Networks Supporting Spatial Attention

**DOI:** 10.64898/2025.12.08.692359

**Authors:** Maria Niedernhuber, David Carmel, Tristan A. Bekinschtein, Corinne A. Bareham

## Abstract

Spatial attention becomes biased toward the right side of space when alertness wanes, but the neural mechanisms underlying this shift remain unresolved. Competing theories propose that spatial attention depends either on a dominant right hemisphere, interhemispheric competition, or interactions between dorsal and ventral attention networks. Here, we used drowsiness as a model to investigate how diminishing alertness reshapes the cortical dynamics supporting auditory spatial attention. Thirty healthy participants completed a lateralised auditory localisation task while high-density EEG was recorded during wakefulness and drowsiness. Behaviourally, drowsiness selectively impaired localisation accuracy for left-sided stimuli, replicating the typical neglect-like rightward attentional bias observed with lower arousal in healthy people, and pathological attention deficits in brain injury patients. Using Dynamic Causal Modelling and Parametric Empirical Bayes, we identified a striking reconfiguration of effective cortico-cortical connectivity across arousal states. During wakefulness, spatial attention was characterised by reciprocal interhemispheric signalling between inferior frontal regions, consistent with balanced bilateral coordination. During drowsiness, this pattern shifted toward reduced right frontoparietal connectivity together with increased left-hemispheric parietofrontal connectivity. These findings suggest that waning alertness destabilises balanced interhemispheric control and allows left-hemispheric processing to dominate attentional processing. Rather than fully supporting existing theories of spatial attention, our results point toward a hybrid mechanism in which neglect-like spatial bias emerges from the interaction between right-hemispheric dysconnection and left-hemispheric dominance under low arousal. Diminishing alertness reshapes interhemispheric coordination rather than simply reducing right-hemisphere function.

## Introduction

Adaptive behavior requires optimal use of spatial attention, the ability to prioritise the processing of sensory input from specific regions in space (Corbetta & Shulman, 2002; Posner, 1980). Although numerous competing theories have been suggested (Bartsch et al., 2023; Brunetti et al., 2008; Chambers, Stokes, & Mattingley, 2004; Clément & Tallon-Baudry, 2025; Corbetta & Shulman, 2011; Desimone & Duncan, 1995; Gaillard & Hamed, 2022; Heilman & Van Den Abell, 1979; Kaiser, Lutzenberger, Preissl, Ackermann, & Birbaumer, 2000; Kinsbourne, 1970; Mesulam, 1981; Shomstein & Gottlieb, 2016; Szczepanski, Konen, & Kastner, 2010; Thiebaut de Schotten et al., 2011; Toba, Migliaccio, Potet, Pradat-Diehl, & Bartolomeo, 2022; Tünçok, Carrasco, & Winawer, 2025; Xia, Chen, Engel, & Moore, 2024; Zuanazzi & Noppeney, 2019), the neural mechanisms of spatial attention remain disputed. An interesting experimental model to adjudicate between competing theories of spatial attention is provided by lateral biases. Typically, healthy and awake individuals exhibit a small attention bias towards the left side of space during visuospatial tasks (pseudoneglect) (Chen et al., 2019; Dickinson & Intraub, 2009; Foulsham, Gray, Nasiopoulos, & Kingstone, 2013; Jewell & McCourt, 2000). However, this attention bias can be reversed under various conditions. Some patients with a stroke in the right hemisphere go on to develop a neuropsychological condition termed hemispatial neglect, characterised by marked inattention to the left side of space (Heilman & Valenstein, 1979; Ting et al., 2011). More subtle rightward attention biases have been observed in neurological and psychiatric conditions such as Parkinson’s Disease (Albert, Bernasconi, Potheegadoo, & Blanke, 2025), or ADHD (Sheppard, Bradshaw, Mattingley, & Lee, 1999). Interestingly, healthy individuals can also experience temporary rightward spatial attention biases (e.g., during vestibular stimulation (Karnath, Himmelbach, & Perenin, 2003)). Numerous studies have documented a transient rightward bias in spatial attention during sleep onset (Bareham, Manly, Pustovaya, Scott, & Bekinschtein, 2014; Chandrakumar et al., 2019; Fimm, Willmes, & Spijkers, 2006; Jagannathan, Bareham, & Bekinschtein, 2022; Manly, Dobler, Dodds, & George, 2005). Similar to clinical neglect (Corbetta & Shulman, 2011; Dietz, Nielsen, Roepstorff, & Garrido, 2021; He et al., 2007), drowsiness is accompanied by a breakdown of frontoparietal connectivity (Goupil & Bekinschtein, 2012; Horovitz et al., 2009; Lacaux, Strauss, Bekinschtein, & Oudiette, 2024; Laufs et al., 2006; Ogilvie, 2001; Sämann et al., 2011). Although several theories explain these neglect-like rightward attentional biases (Bareham et al., 2014; Dobler et al., 2005; Fimm et al., 2006; M. George, Dobler, Nicholls, & Manly, 2005; Manly et al., 2005), they entail distinct predictions about the neural mechanisms which govern them.

According to the right hemisphere dominance theory, the left hemisphere mainly controls attention to the right side of space, whereas the right hemisphere processes information from both left and right spatial fields (Dietz, Friston, Mattingley, Roepstorff, & Garrido, 2014; Heilman & Valenstein, 1979; Heilman & Van Den Abell, 1979; Kaiser et al., 2000; Mesulam, 1981; Thiebaut de Schotten et al., 2011). Therefore, reduced alertness or damage to the right hemisphere leaves only the left to allocate attention, resulting in a systematic rightward bias (Mesulam, 1981; Sadaghiani et al., 2010). At a mechanistic level, the right hemisphere dominance theory of spatial attention predicts disrupted connectivity between early sensory and frontoparietal regions within the right hemisphere when alertness wanes (Dietz et al., 2014; Dietz et al., 2021).

Kinsbourne’s opponent processor theory suggests that inhibition of the ipsilateral cortical hemisphere orients attention toward the opposite side of space (Kinsbourne, 1970). According to this theory, each cerebral hemisphere automatically directs visuospatial attention toward the opposite (contralateral) side of space (the left hemisphere orients to the right, and the right hemisphere orients to the left).The two hemispheres are engaged in a competitive, mutually inhibitory tug-of-war. When one hemisphere becomes highly active, it actively suppresses the orienting mechanisms of the opposite hemisphere. In most individuals, the left hemisphere exerts a slightly stronger orienting drive and inhibition than the right. During drowsiness or neglect then, the right hemisphere no longer inhibits the left, resulting in a bias towards the right side of space (Kinsbourne, 1987). This theory predicts that we will observe an increase in interhemispheric directional connectivity from the right to the left hemisphere (but not vice versa) together with an increase in connectivity within the left hemisphere when alertness wanes (Dietz et al., 2014; Dietz et al., 2021).

Combining aspects of both the right hemisphere and opponent processor theories, the dual network theory states that spatial attention arises from interactions between a dorsal (DAN) and a ventral attention network (VAN) (Corbetta & Shulman, 2002, 2011). The DAN relies on a hemispheric balance, with each hemisphere orienting attention contralaterally (Corbetta & Shulman, 2002, 2011), whereas the right-lateralised VAN is sensitive to novel or salient stimuli. Accordingly, decreasing alertness selectively compromises the VAN, which in turn destabilises the DAN and results in a rightward attentional shift (Dodds, Müller, & Manly, 2009; M. S. George, Mercer, Walker, & Manly, 2008; Malhotra, Parton, Greenwood, & Husain, 2006; Robertson, Mattingley, Rorden, & Driver, 1998). Mechanistically, drowsiness will reduce coupling between regions within the right-hemispheric VAN. This will diminish activity in the connected right-hemispheric DAN but increase connectivity within the left-hemispheric DAN which then dominates spatial attention processing.

Here we set out to disambiguate these theories of spatial attention using reduced alertness as a model. We recorded high-density EEG during an auditory spatial attention task with left and right-sided stimulation when participants were either awake or drowsy. For each state of alertness, we used Dynamic Causal Modelling (DCM) and Parametric Empirical Bayes (PEB) to estimate the context-dependent modulation of effective connectivity induced by left versus right stimulation within a bilateral network spanning auditory temporal, parietal and frontal regions. Rather than a model comparison approach contrasting theories (Dietz et al., 2014, 2014), we applied Bayesian Model Reduction to prune away parameters which do not contribute evidence to the model. Our findings reveal a shift from interhemispheric frontal connectivity in wakefulness to a dominant left frontoparietal network in drowsiness.

## Methods

### Participants

30 healthy right-handed participants (self-reported gender: 20 women and 10 men, age mean = 23.73, SD = 4.88) with normal hearing took part in the study. Handedness was confirmed using the Edinburgh Handedness Inventory (Oldfield, 1971), indicating right-hand dominance across the group (mean Edinburgh score 76.986, SD = 17.884). The mean Epworth sleepiness score (Johns, 1991) for the group was 8.5 (SD = 2.968), indicating normal levels of daytime sleepiness. Participants provided written informed consent prior to the experiment and were reimbursed with NZD $30 worth of cinema vouchers for participation in the study. We initially recruited 36 participants but excluded six EEG datasets from the analysis due to noise. The study was approved by the Victoria University of Wellington Human Ethics Committee committee (approval number 27785).

### Experimental Design

Participants performed a lateralised auditory spatial localisation task under two levels of alertness: awake and drowsy (Figure 2 A). Following Bareham et al. (Bareham et al., 2014), we presented complex harmonic tones at azimuthal positions ranging from -60 to 60 degrees relative to the participants’ egocentric midline. We used guitar chords (D major, D3 at 146.382 Hz) with a duration of 400 ms as auditory stimuli (Bareham, Bekinschtein, Scott, & Manly, 2015) (Figure 2 B). Stimuli were derived from free-field recordings captured via in-ear microphones and reused from previous studies employing this paradigm (Bareham et al., 2014; Jagannathan et al., 2022).

To eliminate non-spatial acoustic cues, the stimulus set comprised both the original recordings and their channel-inverted stereo-flipped counterparts. Stimuli were presented across three azimuthal ranges in each hemispace: near-midline (±0–12 degrees), intermediate (±15–35 degrees), and peripheral (±40–60 degrees). Negative values denote the left hemispace and positive values denote the right hemispace.

First, the participants completed the task while instructed to remain awake and seated upright with eyes closed (awake condition). The stimulus set comprised 100 tones which were balanced across space. Four tones were presented at the midline; the remaining 96 tones were distributed equally between the left and right hemispaces. Specifically, 48 tones were presented in the near-midline range, and 24 tones were each allocated to the intermediate and peripheral ranges. Participants then performed the auditory spatial localisation task while drowsy (drowsy condition). To promote drowsiness, participants switched to a reclined position in a dark, quiet room with eyes closed. They were instructed not to worry about falling asleep as they would be gently roused by the experimenter after three missed trials. In the drowsy condition, 488 tones were presented. 8 tones were presented on the midline and the remaining stimuli were equally divided across both hemispaces; the near-midline range comprised 240 trials whereas the intermediate and peripheral ranges each accounted for 120 trials. Trials were weighted toward the midline to increase task difficulty and evaluate behavioral responses under high perceptual uncertainty during drowsiness, since we did not expect evidence of spatial bias during wakefulness based on previous work (Bareham et al., 2014; Jagannathan et al., 2022).

Regardless of condition, the order in which stimuli were presented was random. To manipulate participants’ temporal expectations of auditory stimulation, each tone was preceded by a variable interstimulus interval. Following previous work (Bareham et al., 2014; Jagannathan et al., 2022), we used interstimulus intervals between 1-2s in the awake condition and 5-8s in the drowsy condition to facilitate the induction of drowsiness. Participants indicated the direction of the tone (either left or right) via a button press with the left and right index finger on the “z” and “m” keys on a keyboard. They were instructed to respond as quickly and accurately as possible. If no response was detected within 5 s of stimulus onset, the trial timed out and the experiment proceeded to the next trial. The task lasted ∼45 minutes depending on response times and omissions. To validate the state of arousal for each trial, EEG data from the 4-s pre-stimulus interval were classified as either belonging to the awake or drowsy condition using the micromeasures algorithm (Jagannathan et al., 2018). We retained only those trials where the algorithmic classification was congruent with the experimental condition. We only included correct trials in our EEG analysis. During wakefulness, we recorded 85.1% correct responses for left-sided stimuli and 89.6% for right-sided stimuli. When participants were drowsy, they gave 78.8% correct responses for left-sided stimuli and 87.5% for right-sided stimuli.

### Behavioural Analysis

Differences in reaction times between wakefulness and drowsiness were assessed using paired-samples t-tests. To investigate whether performance impairments induced by drowsiness are specific to a hemifield, we employed linear mixed-effects models using the lme4 and lmerTest packages in R. Accuracy was modeled with fixed effects of arousal, stimulus laterality and their interaction. We included subject-specific random intercepts to account for the hierarchical structure of the data and inter-individual variability. We determined significance for the fixed effects using likelihood ratio tests comparing full models against nested models excluding the effect of interest. To perform post-hoc comparisons, we employed Kenward-Roger degrees-of-freedom approximations using the emmeans package in R.

### EEG Acquisition and Preprocessing

Continuous EEG data were recorded using a 64-channel BrainVision actiCHamp system with a sampling rate of 500 Hz. EEG data were further preprocessed in Matlab using EEGLAB. Initially, EEG data were downsampled from 500 Hz to 250 Hz and bandpass filtered between 0.5 and 50 Hz. We epoched the data in a time window between -100 to 1000 ms relative to stimulus onset. EEG data were baseline-corrected using a 100 ms pre-stimulus interval. We rejected trials contaminated with noise arising from movements and sweat by visual inspection. To remove physiological noise artefacts, independent component analysis was performed using the EEGLAB Infomax. Components were visually inspected and rejected based on their spatial topography and power spectrum. To prevent classifier bias during multivariate decoding and to ensure equivalent signal-to-noise ratios for PEB/DCM, we equalised trial numbers across the four combinations of arousal state (drowsy/awake) and stimulus laterality (left/right) by randomly deselecting superfluous trials to match the condition with the fewest trials. Having converted EEGLAB to SPM format, we performed source modelling in SPM. EEG sensor positions were projected in 3D space and co-registered to individual structural MRIs. Participants’ MRIs were obtained at the Wellington Hospital prior to the EEG data collection session. Subject-specific forward models were computed using the boundary element method, and the quality of mesh and co-registration was manually verified.

### Decoding

To demonstrate mid-latency differences in auditory neural responses, we employed Time-resolved Multivariate Pattern Analysis (MVPA) for EEG to decode left and right trials in each state of arousal. We down-sampled EEG data to 100 Hz and applied an average mastoid reference (Grootswagers, Wardle, & Carlson, 2017; Trammel, Khodayari, Luck, Traxler, & Swaab, 2023). For each dataset (with at least 25 trials per condition), we trained a decoding pipeline using Z-score normalisation and Linear Discriminant Analysis with automated Ledoit-Wolf shrinkage to discriminate left and right trials within each arousal state. To balance the four classes, we equalised trial counts by randomly discarding trials in the majority classes. This procedure was bootstrapped across 15 iterations per subject to ensure robust estimates.

Classification performance was evaluated using 5-fold stratified cross-validation with five repeats and the Area Under the Curve (AUC-ROC) as a classification performance metric. We smoothed the resulting AUC-ROC time series using a Gaussian kernel (σ=2) to improve the signal-to-noise ratio. Using cluster-based permutation tests, we identified adjacent time points at which stimulus laterality classification performance in each the awake and drowsy condition differed from chance. Cluster-based permutation tests implemented a *Monte Carlo* algorithm with 4096 random partitions and used one-sample two-tailed t-tests for significance testing at α =. 05 (Maris & Oostenveld, 2007). Finally, we reconstructed spatial classifier pattern weights from the multivariate filters using an inverse transform, yielding pattern weights (Haufe et al., 2014). Following previous work (Jagannathan et al., 2022), we compared the resulting spatial activation patterns for the drowsy and awake condition in two time windows (early: 0-400 ms, late: 400-800 ms). Differences between conditions were determined using cluster-based permutation tests with 1000 random partitions based on one-sample two-tailed t-tests at α =. 05 .

### Parametric Empirical Bayes

We used DCM for Event-Related Potentials (ERPs) (David et al., 2006; Kiebel, Garrido, Moran, & Friston, 2008) and PEB in SPM12 (Zeidman, Jafarian, Corbin, et al., 2019; Zeidman, Jafarian, Seghier, et al., 2019) to estimate commonalities and differences in effective connectivity supporting lateral spatial attention between wakefulness and drowsiness. DCM estimates the directed influence that one neural population exerts over another by fitting a generative model to observed neural responses. For that, DCM uses biologically informed neural mass models to simulate voltage amplitude within and between cortical sources. Each source consists of interacting excitatory and inhibitory neuronal subpopulations arranged across canonical cortical layers. Forward, backward, and lateral projections between cortical regions are modeled in accordance with anatomical pathways in the cortical hierarchy. We used DCM to estimate connectivity differences between left and right auditory event-related responses, allowing inference of connectivity changes that best explain observed event-related potentials (David et al., 2006; Friston et al., 2016; Kiebel et al., 2008; Stephan et al., 2010). DCMs contrasting left and right trials were constructed for each participant and arousal state (awake, drowsy) using an ERP model with eight sources in the auditory cortical hierarchy: bilateral A1, STG, IFG and IPC. Sources were modelled as a single equivalent current dipole. We selected eight sources in the ventral frontoparietal pathway based on a previous DCM study contrasting theories of auditory spatial attention (Dietz et al., 2014). Source locations were manually defined based on previous DCM studies of auditory spatial attention and neglect (see Table 1) (Chennu et al., 2016; Dietz et al., 2014; Marta I. Garrido, Kilner, Kiebel, & Friston, 2009; Niedernhuber, Raimondo, Sitt, & Bekinschtein, 2022). Subject-specific head models and forward solutions were used for dipole fitting. We set the number of modes to 8 and the onset to 60 ms. After equalising trial numbers per participant for each of the four folds (left/awake, right/awake, left/drowsy, right/drowsy) by randomly deselecting superfluous trials, models were fitted to EEG data epoched from 0 to 400 ms relative to stimulus onset.

Once individual subject-wise DCMs were estimated for each condition, we applied PEB modelling to characterise group-level effects. PEB is a hierarchical Bayesian Method which treats DCM parameters at the subject-level as inputs to a second-level general linear model. The resulting PEB model can be used to model group-level commonalities and differences in effective connectivity between experimental conditions. We constructed a PEB model over DCM parameters which represent stimulus laterality modulations (left vs right) for each participant and condition. Then, we used PEB to estimate group-level commonalities and differences between awake and drowsy conditions. Using Bayesian Model Reduction, we iteratively tested simplified versions of the parent model by removing expected probability (Ep) parameters with little evidence. Reduced models were then combined using Bayesian Model Averaging (Zeidman, Jafarian, Corbin, et al., 2019; Zeidman, Jafarian, Seghier, et al., 2019), and thresholded at a posterior probability (Pp) ≥ .95.

## Results

### Drowsiness reduces auditory spatial localisation performance in left hemifield

First, we confirmed that our dataset replicates previous findings (Bareham et al., 2014; Jagannathan et al., 2022) of an association between drowsiness and a right-ward bias, whereby the decrease in accuracy that arises with drowsiness is greater for stimuli presented on the left (Figure 2, C, D). Replicating previous studies which identified longer reaction times during drowsiness in this task (Jagannathan et al., 2022), we observed that overall reaction times were prolonged during drowsiness compared to wakefulness (mean difference = 207.51 ms, 95% CI [−291.68, −123.33], t(29)=−5.04, p<.001).

To determine how arousal and stimulus laterality influence performance, we fit linear mixed-effects models on subject-level accuracy with subject-specific random intercepts. Sequential model comparisons via likelihood ratio tests revealed significant main effects for both arousal state (χ^2^_(1)_ = 4. 52, *p* = 0. 034), stimulus laterality (χ^2^_(1)_ = 19. 05, *p* < 0. 001), and an interaction between arousal and laterality (χ^2^_(2)_ = 22. 40, *p* < 0. 001). Post-hoc tests identified degraded accuracy within the left hemifield (*estimate* = 0. 065, *SE* = 0. 024, *t*(93.1) = 2. 75, *p* = 0. 007). In contrast, right-hemifield performance remained stable across states (*estimate* = 0. 015, *SE* = 0. 024, *t*(93.1) = 0. 64, *p* = 0. 521). Consistent with previous work (Bareham et al., 2014; Jagannathan et al., 2022), these findings demonstrate that drowsiness selectively compromises spatial localization accuracy within the left visual field.

We next asked whether drowsiness impaired spatial target detection uniformly, or if the deficit was dependent on stimulus distance. Sequential likelihood ratio tests revealed that adding an interaction term between arousal state and stimulus distance significantly improved model fit (χ^2^_(3)_ = 28. 10, *p* < 0. 001), confirming that behavioural performance reduction in drowsiness is asymmetrically distributed across the visual field. Compared to the awake state, drowsiness was associated with relative reductions in accuracy at 12° (*OR* = 1. 29, *p* < 0. 001), 35° (*OR* = 3. 01, *p* < 0. 001) and 60° (*OR* = 3. 53, *p* < 0. 001). Taken together, the effect of drowsiness becomes stronger the further away from the midline the target appears.

### Drowsiness alters cortical representations of auditory spatial attention

To demonstrate arousal-dependent mid-latency differences in auditory spatial information processing, we compared auditory neural responses to left and right trials using MVPA in wakefulness and drowsiness (Figure 3). Decoding performance was above chance in the awake condition (t = 3.976, p = .0032, time window: 550 - 800 ms) and drowsy condition (t = 6.096, p = .0002, time window: 280 - 800 ms). We also revealed a difference in spatial voltage patterns between the awake and drowsy conditions in a cluster from 250 to 370 ms (t = −535.87,p = .024). No differences emerged in the late time window (400–800 ms).

### Diminishing arousal reconfigures effective connectivity within the auditory attention network

We employed Dynamic Causal Modeling for Event-Related Potentials and Parametric Empirical Bayes to examine how lateralised auditory stimuli modulate effective connectivity in the auditory cortical hierarchy under varying arousal states. Initially, we modelled effective connectivity modulated by spatial location (i.e., the contrast of right versus left trials) within the awake and drowsy states independently, and then we tested for differences between these states. Our analysis revealed arousal-dependent altered connectivity in the auditory cortical hierarchy during drowsiness.

We examined differences in effective connectivity between left and right trials (left > right) during wakefulness and drowsiness. Both the dual network and right hemisphere theories predict that right-hemispheric connectivity will be stronger for left compared to right stimuli during wakefulness, but that this asymmetry will diminish during drowsiness. According to the right hemisphere dominance theory, this asymmetry arises because leftward attention mostly recruits on right hemisphere, whereas rightward attention engages both hemispheres. Similar to the right hemisphere theory, the dual-network model posits this asymmetry since left-sided targets recruit the right VAN for stimulus-driven alerting. However, the dual-network model locates these effects to the VAN (STG, IFG, and IPC), whereas right-hemispheric theory posits that these differences emerge across the entire right-hemispheric hierarchy, including in early sensory areas (A1, STG, IFG, and IPC). According to the opponent processor theory, spatial attention relies on mutually inhibitory interhemispheric interactions. For a left vs right contrast during wakefulness, we will observe increased connectivity across regions in the right hemisphere (A1, STG, IFG, IPC). This will be accompanied by increased inhibition from the right to the left IPC. During drowsiness, mutual inhibition between hemispheres will decrease. Therefore, we expect increased left-hemispheric and decreased right-hemispheric connectivity alongside increased inhibition from the left to the right IPC (Figure 1).

**Figure 1.**
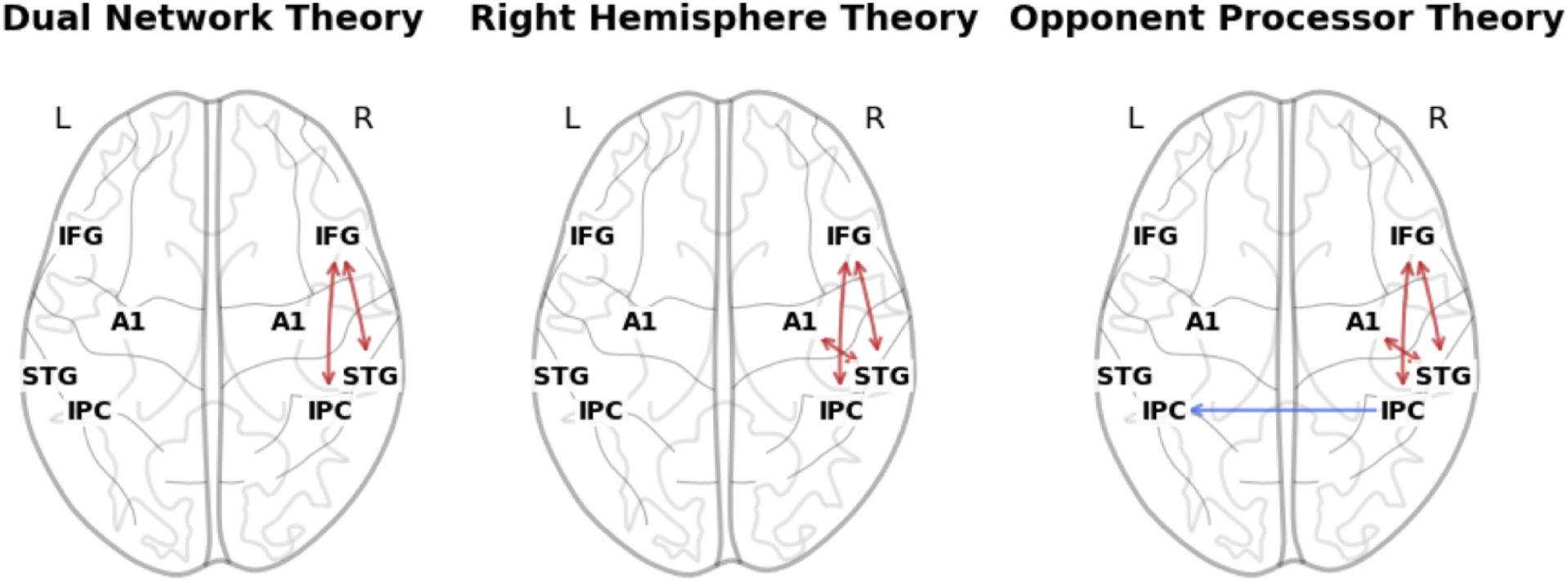
Each theory predicts distinct connectivity modulations for a contrast between left vs right trials in drowsiness. Since these theories were not originally formulated in terms of directed effective connectivity, the predictions shown represent our interpretation of how each account would map onto these connectivity changes within the present DCM framework.

**Figure 2.**
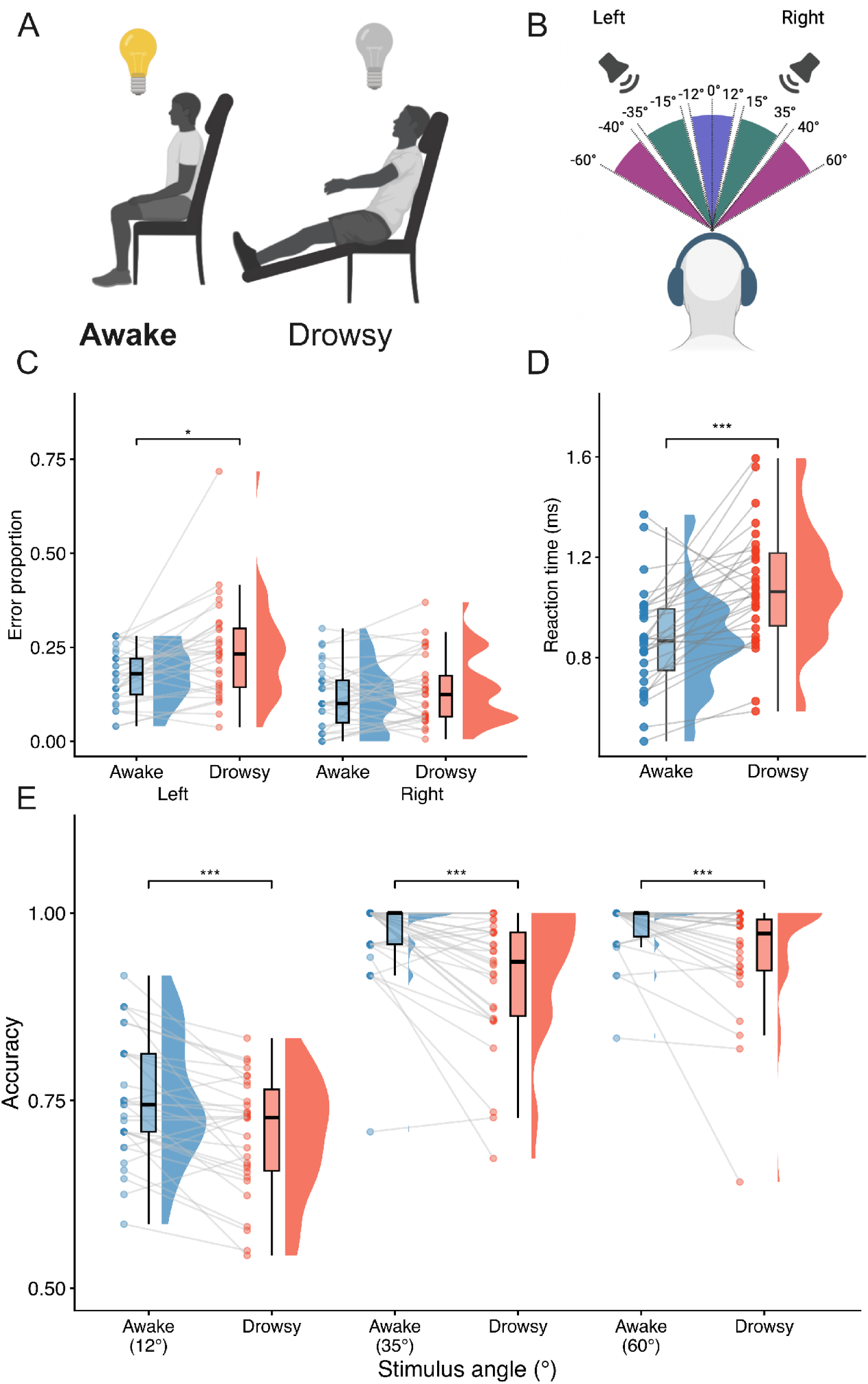
Experimental Design and behavioural performance. (A) Experimental setup. To promote drowsiness, participants remained reclined in a quiet, dark room with their eyes closed throughout the task. During wakefulness, participants sat upright with eyes closed. (B) A schematic depiction of the lateralised auditory spatial localisation task. Participants listened to tones presented from various angles to the left or right of the midline. Spatial locations were categorised into three azimuthal bins per hemispace: near-midline (blue), intermediate (green), and peripheral (purple). (C) Error proportions (1-accuracy) per arousal state and stimulus laterality showing differences between wakefulness and drowsiness for left but not right stimuli. (D) Slower mean reaction times in drowsiness than wakefulness (collapsed across left and right stimuli). Raincloud plots show subject-wise average scores adjacent to a boxplot (indicating median and interquartile range), and kernel density distributions. (E) As expected, overall accuracy (collapsed across left and right stimuli) is highest for the more strongly lateralised tones and gradually reduces towards chance as tones approach midpoint. Raincloud plots show subject-wise average scores adjacent to a boxplot (indicating median and interquartile range), and kernel density distributions. ∗p<.05,∗∗∗p<.001.

**Figure 3.**
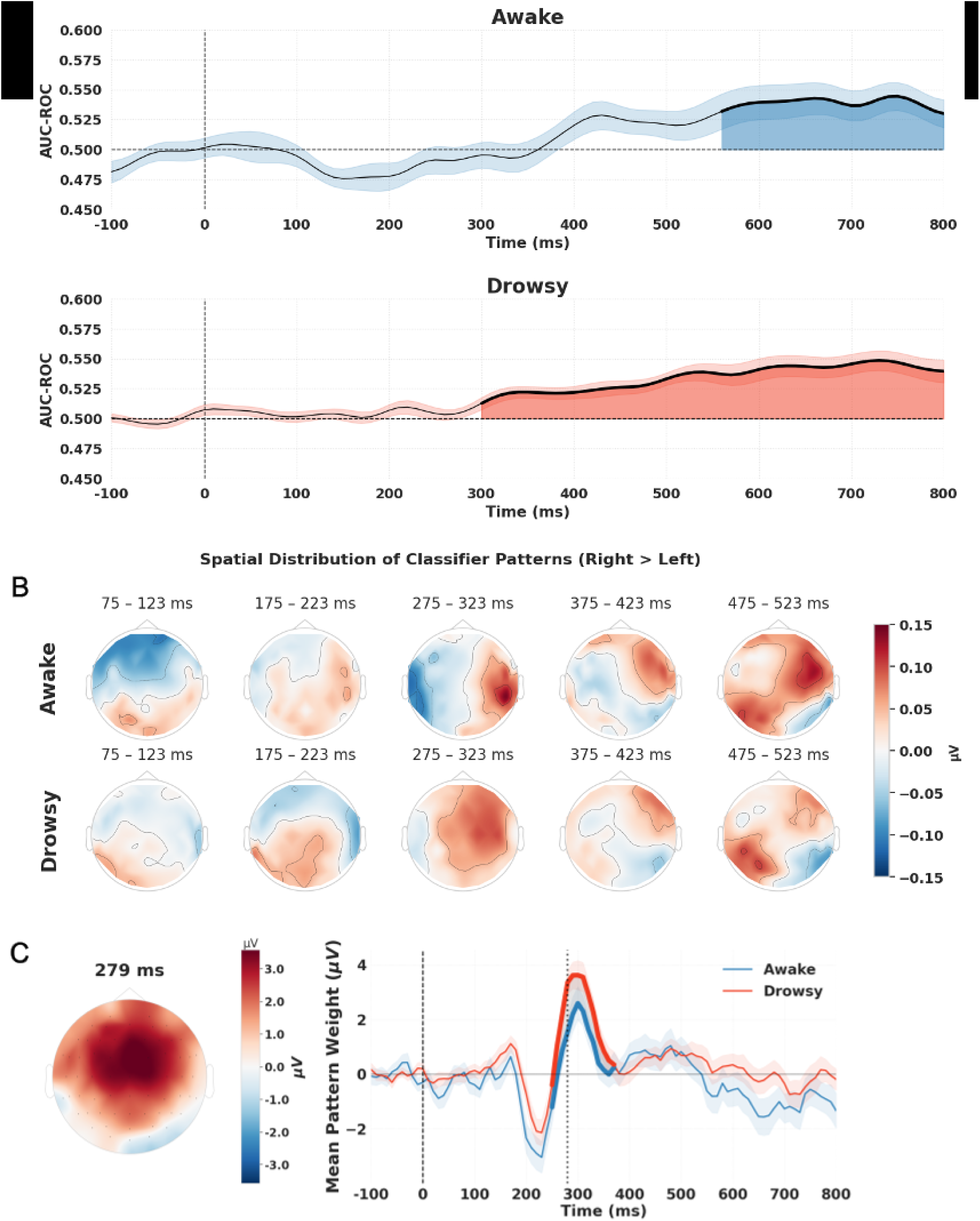
(A) Time-resolved MVPA decoding of stimulus laterality. Mean classifier performance (AUC-ROC) against chance is shown for the awake (red) and drowsy (blue) conditions, with shaded regions indicating the standard error of the mean (SEM). Periods of significant decoding performance are highlighted by thickened lines and shaded underlays. (B) Grand-average scalp topographies for right (R) and left (L) stimuli in both conditions shown in 100 ms intervals post-stimulus. (C) Mean spatial classifier weights pattern time series for the awake and drowsy conditions are plotted with the SEM shaded. Differences between conditions are denoted by thickened lines and grey shading. A vertical dotted line marks the peak difference between conditions. The topographical map showing the difference between the drowsy and awake condition at this time point is displayed on the left.

During wakefulness, we observed increased interhemispheric connectivity between bilateral IFGs when participants processed tones on their left versus their right (depicted using recurrent red arrows in Figure 4, top panel). During drowsiness, we found that connectivity from the right IPC to the left IPC and from the left IPC to the left IFG was increased for left relative to right trials (shown with red arrows between the respective IPCs and left IFG, Figure 4, middle panel). In contrast, connectivity from the right IFG to IPC decreased (displayed with a blue arrow, Figure 4, middle panel).

**Figure 4.**
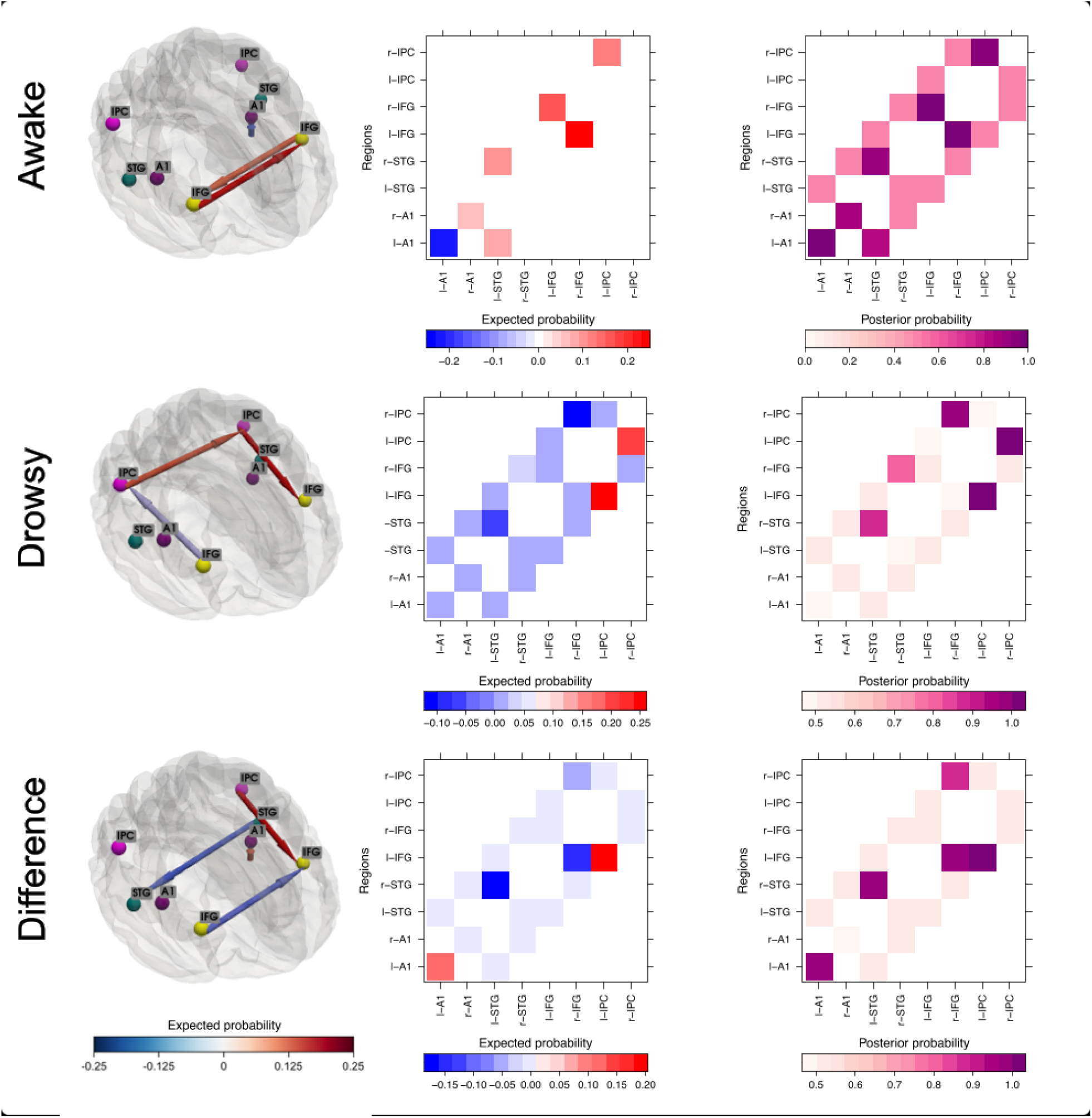
Arousal-dependent modulation of effective connectivity for lateralised auditory attention. PEB results show group-level changes in effective connectivity for the contrast of right versus left auditory trials across different arousal states. PEBs were constructed over DCMs which encoded the contrast between left vs right trials in wakefulness (row 1) and drowsiness (row 2). A PEB-of-PEBs modelled the difference between PEBs for the awake and drowsy states (row 3). Each row depicts effective connectivity modulations between regions in the auditory cortical hierarchy on a glass brain (left), followed by a heatmap of expected probability values (middle) and associated posterior probabilities between connections (right). In the glass brains, arrows indicate modulations of effective connections (B-matrix) between brain areas and their expected probability, plotted on a blue-to-red gradient. In the first two rows, red indicates increasing connection strength between regions when left-sided stimuli are presented (relative to right-sided stimuli), and blue indicates a decrease. In the third row, warm colours indicate that connections are strengthened in drowsiness relative to wakefulness (and cold colours vice versa). All displayed connections are thresholded at a posterior probability of .95.

Finally, we constructed a second-level PEB model contrasting lateralised attention in the awake and drowsy states. When wakefulness was compared to drowsiness directly, we found reduced interhemispheric connectivity from the right to the left IFG (Ep=-0.15, Pp=.98) and from the left to the right STG (Ep=-0.16, Pp=.99) when participants were drowsy vs awake (blue arrows between ROIs, Figure 4, bottom panel). In contrast, connectivity increased from left IPC to IFG (Ep=0.18, Pp>.99) during drowsiness relative to wakefulness (red arrow between ROIs, Figure 4, bottom panel). We also observed more local self-inhibition within the left A1 during drowsiness (Ep=0.13, Pp>.99; red arrow, Figure 4, bottom panel). In a direct comparison between wakefulness and drowsiness, there was positive (but not strong) evidence for a dysconnection from the right IFG to IPC during drowsiness (Ep= -0.06, Pp = 0.88; not shown in plot).

To link behavioural performance and connectivity, we also tested whether accuracy predicts PEB parameters encoding the interaction of arousal and stimulus laterality. We found that the modulatory influence of stimulus laterality on left-right STG connectivity is dependent on an interaction between arousal state and task accuracy (*Ep* = 2. 80, *Pp* = 1. 0). This suggests that a decrease in interhemispheric connectivity between STGs is linked to impaired task performance during drowsiness.

Taken together, our hierarchical analysis of effective connectivity provides a mechanistic account for how diminishing arousal reconfigures connectivity in the auditory cortical hierarchy when spatial attention is engaged. Overall, our results reveal a fundamental shift from a balanced, bilateral network during wakefulness to a left-lateralised network encoding the difference between left and right trials in drowsiness. Rather than the widespread right-lateralised connectivity predicted by the right hemisphere theory, our results showed reciprocal, bilateral interhemispheric signalling between IFGs. We also did not observe the predicted breakdown of connectivity across all levels of the right-hemispheric auditory cortical hierarchy during drowsiness. Instead, we found a reduction in right-hemispheric frontoparietal connectivity during drowsiness, which matches the dual-network theory’s prediction of a compromised right VAN. While the dual network theory predicts left-hemispheric dominance, we identified voltage within the VAN rather than the hypothesised DAN. While wakefulness lacked the unilateral right-to-left inhibition predicted by the opponent processor theory, its prediction for drowsiness aligns with our finding of increased connectivity from the right to the left IPC, as well as from the left IPC to the left IFG.

## Discussion

Our study used a lateralised auditory localisation task across varying levels of alertness to contrast theories of spatial attention. Initially, we replicated behavioural biases in auditory spatial attention during drowsiness (Bareham et al., 2014; Jagannathan et al., 2022), and confirmed differences between neural responses to left-sided and right-sided stimuli in wakefulness and drowsiness. Using temporal decoding, we identified patterns of brain classification weights discriminating left and right auditory stimuli which emerged from ∼280 ms in drowsiness and from ∼550 ms in wakefulness. This finding is consistent with the idea that drowsiness leads to asymmetric neural dynamics between hemispheres in a mid-latency time window, which might underpin the behavioural biases observed when alertness wanes. Using PEB/DCM, we discovered that left-sided stimuli elicit stronger interhemispheric connectivity between inferior frontal and superior temporal regions than right-sided stimuli in wakefulness. Conversely, drowsiness amplified connectivity between frontal and parietal nodes within the left hemisphere, but decreased connectivity in the right hemisphere. Extending previous findings (Bareham et al., 2014), this result suggests that the influence of the left hemisphere in the auditory cortical hierarchy increases as alertness wanes, leading to the characteristic rightward attentional biases observed behaviourally during sleep onset (Bareham et al., 2014; Manly et al., 2005).

Our results do not fully align with any of the theories proposed. For a contrast between left and right trials in wakefulness, the opponent processor theory predicts that the left hemisphere inhibits the right (Kinsbourne, 1970, 1987, 2006), whereas we observed increased interhemispheric flow between IFGs for left compared to right stimuli. For the drowsiness model, our findings comprise elements of both the dual network and opponent processor theories. We identified reduced signaling between the right IFG and IPC for left relative to right trials during drowsiness, which aligns with previous studies showing dysconnection between right frontoparietal regions (Bartolomeo, Thiebaut de Schotten, & Doricchi, 2007; Di Gregorio et al., 2023; Dietz et al., 2014; Dietz et al., 2021; Masina et al., 2022). This observation is congruent with the dual network model’s prediction of dysconnection in the right-hemispheric VAN during drowsiness compared to wakefulness (Corbetta & Shulman, 2011; Dietz et al., 2014).

Simultaneously, we identified increased connectivity from the left IPC to the IFG, coupled with decreased connectivity from the right IFG to the IPC for left compared to right stimuli during drowsiness. This finding supports the opponent processor theory’s prediction of left-hemispheric dominance. While the dual network theory also encompasses the hypothesis that left-hemispheric activity or pattern will increase during neglect, this hyperactivation is thought to be localised in the DAN (Corbetta & Shulman, 2011), and therefore does not align with our observations.

Finally, our connectivity findings are situated within the VAN and do not involve connections between A1 and STG predicted by the right hemisphere dominance theory for the contrast between left and right trials (Dietz et al., 2014; Dietz et al., 2021; Heilman & Van Den Abell, 1979; Kaiser et al., 2000; Mesulam, 1981). We also considered whether our results can be explained by theories of spatial attention beyond these accounts. In 2015, Duecker and Sack inferred a hybrid model of spatial attention that incorporates elements of both the right-hemispheric dominance and the dual-network theory, developed using visual tasks such as line bisection (Duecker & Sack, 2015). The hybrid model of attention emphasises interhemispheric competition involving parietal regions and the frontal eye fields. Since our study uses auditory stimulation, our model did not include the frontal eye field as a region of interest since it is mostly implicated in visual attention tasks (Schall, 2004). In DCM/PEB analysis, connectivity parameters are estimated between a limited number of pre-selected regions, and therefore, our results cannot be used to adjudicate between theories based on predictions about regions not included in the model. Overall, we conclude that the findings presented here do not align with any of the above theories.

Instead, we suggest an alternative structure for connectivity when spatial attention is engaged. Considering the ongoing debate around whether clinical neglect is driven by left-hemisphere hyperactivity or right-hemisphere dysconnection (Bagattini, Mele, Brignani, & Savazzi, 2015; Bartolomeo & Chokron, 1999; Bartolomeo et al., 2007; Brancaccio, Lanza, & Schintu, 2026; Dietz et al., 2014; Dietz et al., 2021; Kaiser et al., 2000; Kaufmann et al., 2024; Koch et al., 2008; Nakamura et al., 2012; Savazzi, Mele, Brignani, & Bagattini, 2013; Thiebaut de Schotten et al., 2011; Toba et al., 2022; Umarova et al., 2011), we propose that both mechanisms might play a role for neglect during drowsiness. Our results indicate that spatial biases during drowsiness emerge from a combination of left-hemispheric decrease in voltage or classification weights and right-hemispheric dysconnection in frontoparietal areas, driven primarily by interhemispheric uncoupling. As alertness decreases, the reciprocal interhemispheric signalling that maintains spatial equilibrium during wakefulness is disrupted. This uncoupling is accompanied by a functional dysconnection within the right frontoparietal network, which concurrently allows the left parietofrontal pathway to become dominant. Consequently, the rightward auditory spatial bias observed in drowsiness is best understood as an emergent network imbalance, demonstrating how interhemispheric decoupling facilitates both right-sided dysconnection and left-sided dominance in frontal and parietal regions.

More recently, predictive coding was used as a framework to explain biased attention in neglect. In previous studies, biased attention in neglect was associated with abnormal precision-weighting of left-sided sensory prediction errors, altered interhemispheric gain control, and disrupted right-hemispheric frontoparietal predictive signalling (Dietz et al., 2021; Doricchi et al., 2021; Marta I. Garrido & Deouell, 2021; Parr & Friston, 2018; Parr, Rees, & Friston, 2018; Vossel et al., 2025). From a predictive processing perspective, our results can be framed as alterations in precision-weighting of sensory prediction errors within the left hemisphere, interhemispheric gain control, and frontoparietal prediction error messaging within the right hemisphere. Within this framework, the behavioral shifts observed during drowsiness reflect a reduction in the precision afforded to left-sided sensory evidence, leading to the selective impairment of targets within the left hemifield. As right-sided influence recedes, this leads to imbalanced interhemispheric gain control, which in turn allows left-hemispheric predictive processes to dominate the attentional hierarchy and bias spatial processing rightward. We note, however, that the predictive coding account can provide a computational framework within which these network-level changes can be interpreted, complementing rather than replacing anatomical theories of spatial attention.

We previously suggested that neglect-like spatial biases during drowsiness might mirror those observed in post-stroke neglect (Bareham et al., 2015, 2014; Manly et al., 2005). In support of this, we found that connectivity from IFG to IPC decreases within the right and increases within the left hemisphere when participants process left-sided stimuli during drowsiness. While dysconnection within the right-hemispheric VAN was previously identified as a neural mechanism for neglect in patients (Dietz et al., 2021; He et al., 2007), there is lively debate around whether the left hemisphere might drive spatial biases in neglect. Various studies supported a view of neglect as driven by right hyperattention, leading to a rightward orienting bias. Theories of post-stroke neglect stipulate that the rightward attention biases seen in neglect patients result from hyperactivity (in PET or fMRI) in the left posterior parietal cortex after a lesion in the right hemisphere (Bartolomeo & Chokron, 1999; Corbetta & Shulman, 2011; Kinsbourne, 2006; Koch et al., 2008; Nakamura et al., 2012). In healthy individuals, parietal transcranial direct current stimulation was found to induce neglect-like spatial bias towards the contralateral hemifield (Geers et al., 2024). However, other groups did not identify a link between left-hemispheric hyperactivity and spatial bias in clinical neglect (Kaufmann et al., 2024; Umarova et al., 2011) or in neuromodulation studies in healthy individuals (Bagattini et al., 2015; Killington, Barr, Loetscher, & Bradnam, 2016). Further work is thus needed to test directly whether similar neural mechanisms govern spatial attention biases in drowsiness and post-stroke neglect, or whether left-hemispheric dominance is specific to spatial bias in drowsiness.

While our findings provide a novel mechanistic account of how alertness modulates spatial attention, it is important to acknowledge that our conclusions rely on EEG-based effective connectivity, which is contingent upon the spatial precision of the specified source model as well as predefined sources. Investigations combining EEG with fMRI data could explore both cortical and subcortical regions that might contribute to spatial attention in wakefulness and drowsiness. Furthermore, our investigation was confined to the auditory modality, and it remains to be seen whether our findings generalise to other sensory modalities such as the visual domain which is primarily affected in neglect patients (Driver & Mattingley, 1998).

In conclusion, our findings provide a mechanistic account of how waning alertness alters cortical networks governing spatial attention. We demonstrate that a rightward spatial attention bias is supported by left-hemispheric dominance and a concomitant reduction in right-hemispheric frontoparietal information messaging. Critically, these results suggest that none of the candidate theories considered in this study fully account for the observed connectivity patterns. Instead, rightward attention biases in drowsiness might be supported by both increased left-hemispheric signaling and right-hemispheric disconnection in frontoparietal regions.

## Code and data

Link to code: https://osf.io/qc9h8/. Link to data: https://doi.org/10.5281/zenodo.19693758

## Funding

This work was funded by a Repatriation Fellowship (1839 RF) and a Small Project Grant (1946 SPG) from the Neurological Foundation of New Zealand, as well as a Marsden Fast-Start Grant (MAU2011) from the Royal Society of New Zealand, awarded to CAB.

## Notes

### Competing Interest Statement

The authors have declared no competing interest.

### Summary of Updates

- Added behavioural analyses, fixed bug in ERP analysis - New title, improved abstract, theoretical framing overhauled - expanded discussion section

